# The role of kinesthetic and visuospatial cues in pain-induced movement avoidance

**DOI:** 10.1101/2023.12.19.572163

**Authors:** Xaver Fuchs, Tobias Heed

## Abstract

**Background:** Avoidance of movements is an important factor in chronic pain. Previous experiments have investigated the involved learning mechanisms by pairing movements with painful stimuli but, usually, other visuospatial cues are concurrently presented during learning. Therefore, participants might primarily avoid these visuospatial rather than the movement-related cues, potentially invalidating related interpretations of pain-induced movement avoidance. Here, we separated kinesthetic from visuospatial cues to investigate their respective contribution to avoidance learning.

**Methods:** Participants used a hand-held robotic manipulandum and, during an acquisition phase, received painful stimuli when performing center-out movements. Pain stimuli could be avoided by choosing a curved rather than direct movement trajectories. To distinguish the contribution of kinesthetic vs. visuospatial cues we used two generalization contexts: either participants executed novel movements passing through the same location at which pain had previously been presented in the acquisition phase; or they executed the same pain-associated movements after having been reseated, so that the hand did not pass through the pain-associated location.

**Results:** Avoidance generalization was comparable in both contexts, and remarkably, highly correlated between them. Our findings suggest that both visuospatial and kinesthetic cues available during acquisition were associated with pain and led to avoidance.

**Conclusions:** Our research corroborates the fear-avoidance pain model and previous studies’ findings that pain can become associated with movements. However, our study indicates that visuospatial cues also play a critical role. Future studies should distinguish movement-related and space-related associations in pain learning.

**Significance:** Chronic pain is a significant health issue typically attributed to maladaptive learning of pain-movement associations and movement avoidance. We demonstrate that visual cues can play a similarly important role as movement cues in pain learning. This aspect has not previously been considered and has likely confounded previous research findings.

## 1 Introduction

Chronic pain is characterized by pain persisting without a clear physical cause. Today, chronic pain is considered one of the leading health problems worldwide, affecting as many as 19% of adults according to a survey conducted in 15 European countries and Israel (Breivik et al., 2006).

We still lack a comprehensive mechanistic understanding of the development of chronic pain, which makes it difficult to develop effective treatments. Pain chronification is a complex process that involves somatic, psychological, and social factors (Flor & Turk, 2011). A common notion is that initially acute pain becomes chronic through the mediation of learning and memory (Apkarian et al., 2013; Flor & Turk, 2011; Mansour et al., 2013) in interaction with other cognitive and emotional variables (Fuchs, Bekrater-Bodmann, et al., 2015). The “fear-avoidance model” of pain captures the idea that cognitive and learning processes play a pivotal role in pain chronification (Lethem et al., 1983; Vlaeyen & Linton, 2000). The model posits that chronic pain starts with an incident in which a painful stimulus (the unconditioned stimulus; UCS) causes a defensive response as an unconditioned response (UCR) and, via classical conditioning, the UCS becomes associated with an initially neutral stimulus (NS) that is present during the occurrence of the UCS. Over time, the NS becomes a conditioned stimulus (CS+) that subsequently elicits fear as a conditioned response (CR). As a concrete example, back pain may initially have been caused by bending over. Thus, over time the bending-over movement becomes the CS+ and elicits fear of pain. Moreover, when the patient avoids the bending-over movement (i.e., the CS+), no pain occurs, supporting learning of the avoidance response via negative reinforcement—a classical example of operant conditioning. Unfortunately, the prolonged physical inactivity resulting from such movement avoidance as well as avoiding daily activities that may involve the feared movement, both have detrimental effects on musculoskeletal and cardiovascular health and can, therefore, aggravate pain problems. For many patients, these accumulating and persistent difficulties result in withdrawal from physical and, eventually, also social activities; this behavior, in turn, promotes negative mood, which can reduce pain tolerance. In this manner, affected persons end up in a vicious, self-amplifying circle of fear, avoidance, negative mood, and pain.

Evidence for the negative consequences of pain-related fear for pain and disability comes from questionnaire studies in pain patients (Crombez et al., 1999; Gheldof et al., 2010; Picavet et al., 2002). However, the characterization of the interrelationships proposed in the model would benefit from dedicated, experimental manipulation of specific pain and movement aspects. Indeed, a series of studies has taken this approach and induced pain-related movement avoidance in healthy participants (Glogan et al., 2020, 2022; Karos et al., 2017; Meulders et al., 2011, 2016). For instance, one study investigated the acquisition of avoidance behavior (Meulders et al., 2016) by asking healthy participants to use a robotic arm to steer a visually presented, virtual ball from a fixed start to a target position. The movement had to proceed through one of three adjacent visually presented virtual gates halfway along the path that were arranged such that there were three possible movement trajectories: a direct path, a slightly curved, and a strongly curved path. The robotic arm penalized curved trajectories through a counterforce, so that they required more physical effort than the direct, straight trajectory. For the experimental group, the probability of receiving an electrical pain stimulus was 100% for the direct trajectory and lower for the curved trajectories, effectively resulting in a trade-off between the probability to experience pain and an effort to attempt its avoidance. In a yoked control group, participants received the same pain stimuli as the experimental group but without a systematic relationship between pain and movement trajectory. Only the experimental group acquired higher levels of pain expectation and fear of direct trajectories and avoided the direct more than the curved trajectories. This result suggested avoidance learning in the experimental group. A follow-up study then introduced a generalization manipulation (Glogan et al., 2020): After training with the three gates, participants had to move through three new gates and never received any pain stimulation. The experimental group reported higher pain expectancy and fear related to new movement trajectories that were spatially close to that trajectory which had been associated with pain in the fear-learning scene. In contrast, avoidance behavior did not generalize to the new trajectories.

These experimental results imply, thus, that fear but not movement avoidance generalize. However, several confounds potentially limit the interpretation that the participants had, in fact, learned avoidance of *movements* in the initial training phase. This is because the paradigm of these studies provided visual, proprioceptive, and motor cues all at once; it is, therefore, unclear which information was associated with pain.

Maybe most prominently, it is possible that participants rather avoided a *location in space* rather than a specific movement. This is possible because both the virtual ball that participants guided, and the gates through which the ball had to pass were presented visually and accordingly always occurred at a fixed location in space. Thus, even if choosing a particular gate resulted in one of three distinct movement trajectories, it is unknown whether any movement-related information was ever relevant to participants developing fear and avoidance. Alternatively, they may have learned to fear and avoid that the ball is seen in the location of the gate associated with pain stimulation.

These various possible explanations map to very different kinds of real-life analogies. To give an example, imagine picking a single flower from a thorny rosebush. As the hand reaches the rose, it touches a thorn that painfully stings the hand, and you pull back. What will you associate with this painful experience and, accordingly, do next? According to the fear-avoidance model, the pain is associated with the movement. Thus, you would avoid grasping the rose again in the same way, using the same movement path, but not necessarily grasping the same rose all together. An alternative possibility is that the pain is associated with the location of the rose: the one you had chosen turns out to be surrounded by many thorny stems, and so it may be sensible to avoid this particular location in space. In this case one would avoid grasping for the rose, irrespective of the grasping movement. Our point is that in analogy with this example, the cited paradigms (Glogan et al., 2020; Meulders et al., 2016) might foster a learning process that resembles the second alternative. Figuratively speaking, participants may avoid a gate in the discussed experiments like they might a stingy area of the rosebush – not because the movement hurts, but because pain occurs along the movement’s spatial trajectory or at its target location. Given these rather distinct possibilities of the potential causes of pain learning associated with movement and visual cues, it appears desirable to experimentally dissociate movement and spatial location.

We addressed this differentiation in an experiment in which participants either made identical reaching movements in different portions of the workspace or made distinct movements that crossed through a common spatial region of the workspace. This experimental strategy allowed us to dissociate the factors movement and space.

## 2 Methods

### 2.1 Participants

The study comprised two groups, an experimental (“Conditioning”) group and a matched comparison (“Yoke Control”) group. We aimed for a sample size of 30 participants per group—an estimation that is well in the range of participants used in previous studies (Meulders et al. (2016): 25 participants per group; Glogan et al. (2020): 32 participants per group). We did not perform an a-priori power analysis due to absence of prior data for our paradigm that could have served as an effect size estimate. However, we performed a post-hoc sensitivity analysis (see section 3.3.1) as is recommended to gauge power post-hoc (Lakens, 2022). We excluded the data of two participants due to technical errors during acquisition. The *Conditioning* group therefore consisted of 28 participants (13 female based on self-report; mean age 28.21 ± 8.79 years). We recruited a *Yoke Control* group of 28 participants that were each matched with a participant from the *Conditioning* group based on sex (13 females; mean age 27 ± 8.75 years; range 18-54 years). We also matched groups based on age. For 21 cases the age difference between the matched participants was less than 2 years; for 7 participants it was between 3 and 5 years. All participants were right-handed as assessed by the Edinburgh Handedness Inventory (Oldfield, 1971). After screening the movement data, one additional participant from the *Yoke Control* group had to be excluded (see section 3.3.12), resulting in an analyzed data set of 28 participants in the *Conditioning* group and 27 in the *Yoke Control* group.

Participants were recruited through advertisement boards at Bielefeld University and by direct contact. They received course credit for participation. Potential participants were not tested if they reported neurological or mental disorders, severe physical or sensory impairments, chronic pain, or use of medication that affects mood, cognition, or pain sensitivity. The study was approved by the ethics committee of Bielefeld University and participants gave written informed consent.

### 2.2 Procedures

The session comprised the assessment of pain thresholds as well as a movement task during which participants occasionally received painful electrical stimuli. Participants were informed about the procedures. They were informed that the intensity of the stimuli presented during the movement task was adjusted to their personal pain threshold and about the possibility to down-regulate the intensity or terminate testing at any time. The session lasted about one hour.

### 2.3 Electrical pain stimuli and assessment of pain thresholds

Two cup electrodes (Ag/AgCl) were attached to the participants’ dorsal right wrist to apply electrocutaneous stimuli. They were filled with Abralyt 2000 conductive gel and attached to the skin with medical adhesive tape. Electrical stimuli were generated by a bipolar constant current stimulator (Digitimer DS5, Digitimer, Welwyn Garden City, UK) set to an output current of 50 mA. Participants’ pain thresholds and pain tolerance thresholds were assessed with a custom Matlab program (MATLAB R2015a, Mathworks, Natick, Massachusetts, USA) that controlled the Digitimer via a USB-connected digital-analogue-converter (NI-USB 6343, National Instruments, Austin, TX, USA). For the assessment of both pain intensity and pain tolerance thresholds, the program presented three ascending pulse series (pulse duration: 15 ms), respectively, that started with an output intensity of 1% of the Digitimer’s maximum output energy and then incrementally increased stimulus intensity in 1%-steps at a frequency of 0.5 Hz. For pain threshold assessment, participants had to press the space key as soon as the electrical stimuli felt painful; for pain tolerance thresholds, they pressed the key when they felt that they could not tolerate any further intensity increase. This latter key press immediately ended the presentation of electrical stimuli. The stimulus intensities present at the time of the key press were averaged across the three series to estimate the individual’s pain and pain tolerance thresholds. In keeping with the above-described prior studies (e.g. Meulders et al., 2016), we aimed to use stimuli in the painful range that require some effort to tolerate for our conditioning experiment (see below, 2.2.2). To this end, we used the intensity at 50% between the pain threshold and the pain tolerance threshold for each participant (Lang et al., 2009).

#### 2.3.1 Movement Task

Participants operated the handle of a robotic manipulandum (KINARM End-Point Lab, Kinarm, Kingston, Ontario, Canada) with their right hand to perform diagonal reaching movements. The Kinarm is a device that allows recording handle position as well as the forces applied on the handle by the hand, both at a rate of 1000 Hz. The device projects visual stimuli from a top-mounted horizontal monitor onto a mirror, so that cursor and visual stimuli appear in the same plane as the handle. In our setup, vision of the handle and arm/hand was prevented. Instead, hand position was projected as a 0.5 cm round, white cursor onto the mirror. The Kinarm can apply forces to the handle to create virtual force channels that constrain movement to a predefined trajectory. In some experimental conditions (see below), we manipulated the position of the seat on which participants sat. To this end, the chair was installed on a rail that allowed positioning the chair at two positions 20 cm apart on the left-right axis.

At trial start, participants saw a starting position (red circle with 1 cm diameter) and moved the hand cursor there. When the cursor had been fully within the start circle for 500 ms, the start position disappeared. 1000 ms later, a target position (red circle with 1 cm diameter) appeared, and participants moved the cursor to the target position. The target position then flashed green and disappeared, which ended the trial. The target position was always 20 cm away in depth and either 20 cm to the left or right from the starting position, resulting in 28.28 cm from start to target position (see Fig. 1).

**Figure 1.**
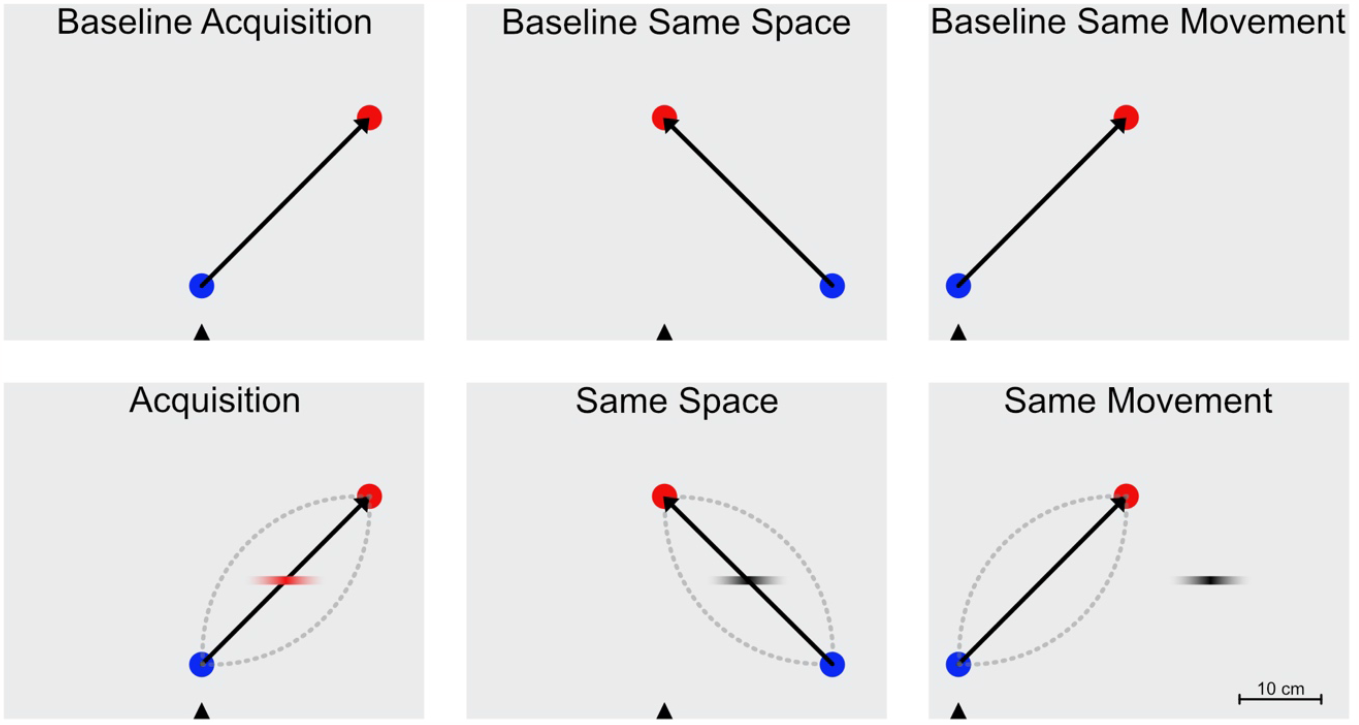
Experimental phases. The experiment began with three baseline phases (upper row) that matched the phases testing for generalization of fear. Participants did not receive painful stimuli during this part of the experiment. Avoidance acquisition and generalization involved 3 phases (lower row): In the *Acquisition* phase participants received painful stimuli. The *Conditioning* group always received pain stimuli when crossing through an invisible, horizontal trigger line positioned halfway along the movement path. The gradient of the red line signifies varying pain intensities depending on the exact location at which the trigger line was crossed. Importantly, the *Conditioning* group could avoid the pain by using curved movements, illustrated by the dotted lines; the *Yoke Control* group received pain stimuli in random trials, irrespective of their movement behavior. Two conditions without pain stimuli tested how the learned avoidance behavior was associated with space and was maintained when performing a different movement that crossed the same space, or, when the same movement was performed in a different area of the workspace, achieved by changing the participants’ sitting position 20 cm to the left, illustrated by the black triangle. In the *Same Space* and *Same Movement* phases, no pain stimuli were presented; the black gradient in the figures indicates the position where the painful stimulus had been presented in the *Acquisition* phase.

Before participants started the movement task, they were informed that they would perform reaches from a start to a target position. Moreover, they were told that “whether or not they would receive pain depended on their behavior”; in the context of the written instructions, it was clear that “behavior” referred to motor behavior. Participants were then familiarized with the Kinarm setup and given the opportunity to practice controlling the cursor with the handle and explore the operating space.

The session comprised 9 phases that were separated by short breaks in which participants rated their level of fear of movement on a numerical rating scale (NRS) ranging from 0 (no fear) to 10 (strongest fear), whether they had received pain stimuli (yes/no), and, in case they had received pain stimuli, how intense the pain was on an NRS from 0 (no pain) to 10 (strongest pain).

The 9 phases consisted of 3 baseline phases at the beginning of the session that were presented in randomized order, followed by 3 identical acquisition blocks, followed by 2 generalization phases, again in randomized order, and, lastly, a forced extinction phase. Before each baseline phase, participants were explicitly told that no pain stimuli would occur in the upcoming phase and that they should move normally. The cable connecting the Digitimer and the electrodes was disconnected before the eyes of the participant to emphasize credibility of this instruction and reduce any potential fear related to pain expectation. The phases were the following (for an illustration, see Fig. 1):

- *Baseline Acquisition (25 trials)*. right-side sitting position and outwards movements, from the center, aligned with the body-midline, towards the far-right target.
- *Baseline Same Space (25 trials)*: right-side sitting position and movements inwards, from 20 cm right of the body-midline towards a center-far target position.
- *Baseline Same Movement (25 trials)*: left-side sitting position and movements like in the *Baseline Acquisition* (and *Acquisition*, see below), from the center to a far-right target.
- *Acquisition (3 blocks with 20 trials each)*: Participants were told that, from now on, they might receive painful stimuli and the electrode cable was connected to the stimulator. The movement targets were identical to those of *Baseline Acquisition*. When the handle crossed an invisible, virtual trigger line positioned halfway along the diagonal between start and target (10 cm width, i.e., 5 cm to each the left and right of the midpoint; see Fig. 1) an individually adjusted pain stimulus was triggered. To reinforce curved avoidance movements, stimulus intensity was at the full intensity when participants crossed the line in the middle, i.e., when performing a straight trajectory between start and end position; pain intensity decreased towards the outside of the line, following a Gaussian density function with a standard deviation of 1.67 cm from the midpoint towards each side (see Fig. 1).
- *Same Space (20 trials)* generalization. Participants were not informed about any changes regarding the pain stimulus, but no painful stimuli were presented any more. Start and end positions were identical to *Baseline Same Space*; note that the movements directly crossed through the line that triggered the painful stimulus in the *Acquisition* phases. This phase assesses how avoidance of straight trajectories is associated with space and therefore generalizes to new movements crossing through a common spatial location.
- *Same Movement (20 trials)* generalization. Again, no pain stimuli were presented but again this change was unannounced. The sitting position was changed to the left and the start and end points instructing the required movement were identical to those of *Baseline Same Movement*. Note that it is also the same movement as in *Acquisition*, starting at the body midline and then moving towards a target to the top right. The new sitting position, however, had moved the body to a different portion of the workspace. Thus, although the performed movement was identical to before, it did not cross the spatial area at which the painful stimuli had been presented in the *Acquisition* phase). This condition assesses how avoidance of straight trajectories is associated with the movement proper and therefore generalizes to a new portion of space that is crossed with an identical movement.
- *Forced Extinction (10 trials)*. The movements were exactly like in the *Acquisition* phase. As in the baseline phases, the electrodes were disconnected, and the participants were told that no pain will occur. The movement was constrained by a force channel so that participants were coerced to perform straight movements crossing the midpoint of the line. This phase was merely included to “wash out” the learned avoidance behavior but was not included in any of the analyses.

The *Yoke Control* group underwent the same phases as the *Conditioning* group, but the contingency in the *Acquisition* phase was absent so that no systematic relationship between movement and avoidance of pain could be learned. For each participant of the *Yoke Control* group, pain stimulation was matched to that of one participant in the *Conditioning* group of the same sex and similar age (see above, section 2.1.); stimulation was matched with respect to the trial in which the match had been stimulated, as well as the intensity modulation related to how far the match’s movement had been away from the trigger line’s midpoint; however, intensity was set relative to the yoked participant’s individual pain and intensity thresholds. As a concrete example, if a participant of the *Conditioning* group performed a slightly curved trajectory in trial 5 of the second *Acquisition* phase and, accordingly, received a pain stimulus at 80% stimulus intensity, then the respective yoked participant also received a pain stimulus at 80% of his/her determined stimulus intensity in exactly that trial and when his/her trajectory crossed the midline in depth between the start and end points, no matter how s/he executed the movement in that trial. As a result, the *Yoke Control* group received the same number of pain stimuli and at comparable (individual) intensities as the *Conditioning* group, but pain was independent of their movement behavior.

### 2.4 Data analysis and statistics

#### 2.4.1 Sensitivity power analysis

We carried out two sensitivity analyses; one for the sensitivity for discovering that the change from baseline to acquisition is significantly larger in the *Conditioning* than the *Yoke Control* group – i.e., the online learning effect –, and one for discovering that the change from baseline to the generalization phases is larger in the *Conditioning* than the *Yoke Control* group – i.e., the fear generalization effect. Both sensitivity analyses revealed that our design was sensitive enough to detect effects of half the size observed in our study with a power of 0.9 suggesting that the study was well powered (see the supplementary material).

#### 2.4.2 Data screening and exclusion of movement data

We first removed very slow trajectories using a criterion of exceeding 3000 ms to reach the target, because such long reach times indicate that participants did not perform a regular, instructed reach to the target. This was the case for 334 trials (3.4%). Most instances were caused by a single participant from the *Yoke Control* group for whom this applied in 127 trials (73% of this participant’s data). We therefore removed this participant’s dataset from statistical analysis.

#### 2.4.3 Spatial preprocessing and operationalization of avoidance as trajectory curvature

We operationalized avoidance behavior through assessment of trajectory curvature. Recall that we instructed participants that the occurrence of pain stimuli depended on their movements. The rationale, therefore, is that when no fear to experience pain exists, then the movement will be straight between start and end point. Of course, given skeletal and muscular constraints, movements may naturally deviate from a straight line. Therefore, we assessed trajectory curvature in separate baseline conditions for each experimental movement; and computed avoidance as the difference in curvature between the respective condition and the baseline conditions in which participants performed identical movements (in an identical space).

We computed curvature by first reducing trajectories to 100 equidistant points using the “spatialize” function from the *mousetrap* package (Wulff et al., 2021) for the R programming language (R Core Team, 2023). After this step, each trajectory had the same number (100) of x/y coordinates, which facilitates comparisons of their shapes. We then computed the deviation, that is, the Euclidian distance from the straight line between the start and the target for each of the 100 points and averaged these deviation values over the 100 points to yield a per-trial average distance from the straight line. This distance is closely related to the area under the curve, a standard index for curvature in movement trajectories (Wulff et al., 2021).

#### 2.4.4 Statistical analysis of movement trajectories

We entered curvature values as the dependent measure into our statistical analyses. We used linear mixed models (LMM) to compare the degree of trajectory curvature between groups and conditions. Inspection of the LMMs’ residuals revealed deviation from a normal distribution, which we remedied by a log-transformation of the curvature values. The log-transformation improved the fit of the LMMs and made the derived statistics more credible but did not affect the qualitative interpretation of the results (see the supplementary information).

We fitted LMMs to address the following questions:

1. One LMM served as manipulation check of acquired avoidance behavior by testing whether the trajectory curvature increased in the *Acquisition* as compared to the three baseline conditions, and whether this increase was stronger in the *Conditioning* as compared to the *Yoke Control* group. We included two fixed factors: *Group* (two levels: *Conditioning* and *Yoke Control*), and *Phase* (4 levels: *Baseline Acquisition, Baseline Same Space, Baseline Same Movement*, and *Acquisition*). Separating the three baseline conditions was important because they comprised different movements, so that movement curvature could differ between them. In contrast, the factor level “*Acquisition*” combined the three blocks of the *Acquisition* phase because they were all identical. The hypothesis that curvature increases in *Acquisition* as compared to the *Baseline* conditions especially for the *Conditioning* group is captured by the *Group* × *Phase* interaction.
2. A second LMM compared avoidance between conditions that tested movement vs. spatial association by comparing the differences in trajectory curvature of the movement and spatial generalization against their respective baseline phases (i.e., *Same Space* vs. *Baseline Same Space*, and *Same Movement* against *Baseline Same Movement*). We included three fixed factors: *Group* (two levels: *Conditioning* and *Yoke Control*), *Phase* (two levels: *Baseline* and *Generalization*), *Association* (two levels: *Space* and *Movement*) and all their interactions. We hypothesized a larger difference between *Generalization* and *Baseline* for the *Conditioning* group, which is expressed by the *Group* × *Phase* interaction. If avoidance of straight trajectories is more strongly associated with space than with movement, or vice versa, this would be indicated by a significant *Group* × *Phase* × *Association* interaction.

For all LMMs, including those described in the following sections, we followed the recommendation to use the most complex random effects structure that does not result in failure of model convergence (Barr et al., 2013). We followed up on significant effects of the highest rank (interaction term in case of significance, main effects in case of significance and absence of interactions) by computing pairwise comparisons between estimated marginal means (EMMs). We restricted the tested comparisons to the contrasts of interest. We corrected p-values for multiple testing using the false discovery rate (Benjamini & Hochberg, 1995).

To analyze whether participants’ avoidance of the affected portion of space vs. avoidance of a specific movement were associated with each other, we computed Pearson correlations of the curvature values between the *Same Space* and *Same Movement* conditions. A positive correlation indicates that participants who avoid based on space also avoid based on movement; a negative correlation indicates that participants either avoid based on space *or* movement, and the absence of a correlation indicates that avoiding space is independent from avoiding movements.

#### 2.4.5 Statistical analysis of fear and pain ratings

We analyzed whether the reported fear of movements differed across phases and whether this difference depended on the group. The LMM included two fixed factors: *Group* (two levels: *Conditioning* and *Yoke Controls*), and *Phase* (6 levels: *Baseline Acquisition, Baseline Same Space, Baseline Same Movement, Acquisition, Same Space*, and *Same Movement*). As before, the factor level “*Acquisition*” combined the three identical *Acquisition* blocks.

Lastly, we tested whether the *Conditioning* and the *Yoke Control* group perceived pain intensity differently and whether ratings differed between the three *Acquisition* blocks. We computed an LMM with two fixed factors: *Group* with (two levels: *Conditioning* and *Yoke Control*), and *Block* with (three levels: Acquisition 1, 2, and 3). Because the frequency of pain stimuli and stimulation intensity could differ across blocks depending on how strongly participants curved their trajectories, we included the average stimulus intensity (percentage of the full intensity stimulus) as a covariate.

#### 2.4.6 Software

For all analysis, we used R (R Core Team, 2023). We computed LMMs with the packages *afex* (Singmann et al., 2023) and *lme4* (Bates et al., 2015). EMMs and post-hoc comparisons were computed using the *emmeans* package (Lenth et al., 2019). Spatial preprocessing of the movements trajectories was done using *mousetrap* (Wulff et al., 2023). Results figures were created with *ggplot2* (Wickham, 2009).

#### 2.4.7 Availability of data and code

All data and code to reproduce the statistical results and figures presented in this article are publicly available on the website of the Open Science Framework accessible via the following link: https://osf.io/rjz36/.

## 3 Results

### 3.1 Acquisition of avoidance behavior

As expected, trajectories were nearly straight in the baseline phases but curved in *Acquisition* phase (see Figure 2).

**Figure 2.**
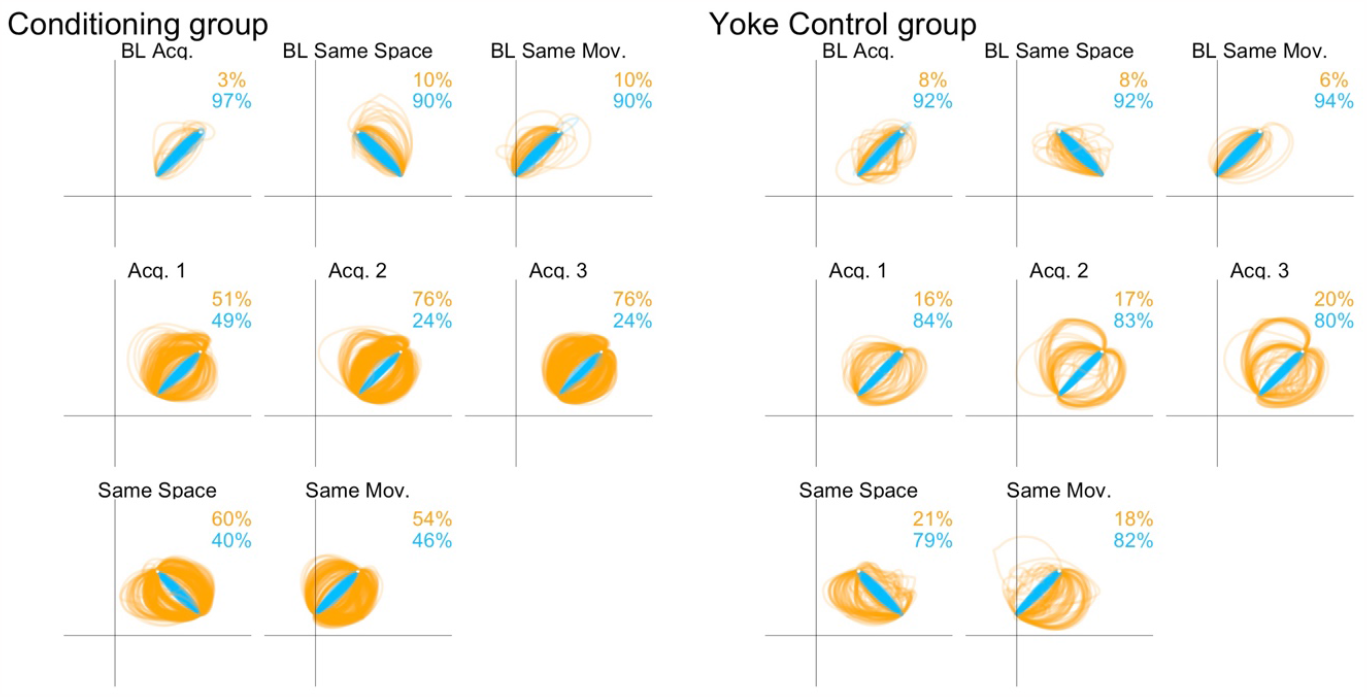
Movement trajectories of the *Conditioning* (left) and *Yoke Control* group (right) in the eight phases of the experiment. In each trial, participants reached from the start (close to the x axis) to the target position. In the figures, all trajectories of all participants are overlayed. The colors, blue and orange, highlight straight and curved trajectories, respectively, and their proportions are given by the percentage numbers. For illustration purpose, we classified the trajectories into straight and curved by first calculating the curvature values for each trial (average distance from the straight path, see text for details) and then coding all trials below the average curvature value as “straight” and all above as “curved”. The proportion of curved movements increased in the *Acquisition* as compared to the baseline conditions and was also elevated compared to the baseline conditions when the pain stimuli were omitted in the generalization conditions (*Same Space* and *Same Movement*).

The LMM performed as manipulation check showed a significant *Group* × *Phase* interaction (F(3, 53) = 10.1, *p* < 0.001; for other effects see Table 1) indicating that the change in curvature over conditions differed between the *Conditioning* and *Yoke Control* groups. This interaction was followed up by statistical comparison of the EMMs, revealing that curvature during the *Avoidance* phase was stronger than in the three baseline phases in the *Conditioning* group but not in the *Yoke Control* group (see Table 2 for all relevant statistical comparisons between conditions).

**Table 1:** Results of the linear mixed model analyzing the increase in trajectory curvature from baseline to *Acquisition* phases between the groups.

| Term | num DF | den DF | F | p | sign. |
| --- | --- | --- | --- | --- | --- |
| Group | 1 | 53 | 8.0 | 0.007 | ** |
| Phase | 3 | 53 | 23.7 | <0.001 | *** |
| Group × Phase | 3 | 53 | 10.1 | <0.001 | *** |

**Table 2:** Post-hoc comparisons between the *Acquisition* and the baseline phases for the groups. BL=Baseline; units are log cm.

| Contrast | Group | Difference | DF | t | p | sign. |
| --- | --- | --- | --- | --- | --- | --- |
| BL Acq. - Acq. | Conditioning | -1.9 | 53.0 | -9.4 | <0.001 | *** |
| BL Same Space - Acq. | Conditioning | -1.5 | 53.0 | -7.1 | <0.001 | *** |
| BL Same Mov. - Acq. | Conditioning | -1.6 | 53.0 | -8.1 | <0.001 | *** |
| BL Acq. - Acq. | Yoke Control | -0.3 | 53.0 | -1.6 | 0.165 |  |
| BL Same Space - Acq. | Yoke Control | 0.0 | 52.9 | -0.2 | 0.833 |  |
| BL Same Mov. - Acq. | Yoke Control | -0.3 | 53.0 | -1.3 | 0.246 |  |

There was no statistically significant difference between the groups in any of the baseline phases (see Table 3 for all relevant statistical comparisons between the groups). In contrast, curvature was significantly higher in the *Conditioning* than in the *Yoke Control* group during the *Acquisition* phase, (Difference = 1.5 log cm, t(53) = 5.1, p < 0.001; see Figure 3A). In sum, the introduction of pain stimuli affected trajectory curvature – indicative of stronger avoidance – most strongly in the *Conditioning* group, that is, the group in which pain stimuli depended on behavior, confirming that our experimental manipulation affected the two groups as intended.

**Table 3:**
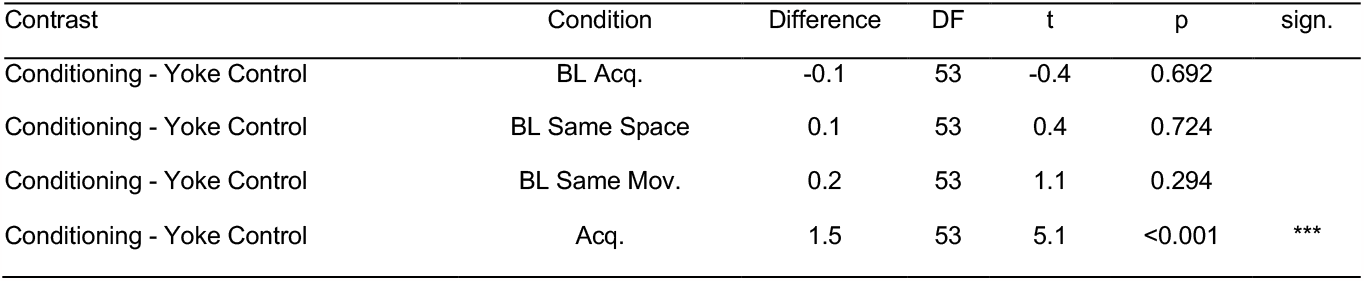
Post-hoc comparisons between the groups in the *Acquisition* and the baseline phases. BL=Baseline; units are log cm.

| Contrast | Condition | Difference | DF | t | p | sign. |
| --- | --- | --- | --- | --- | --- | --- |
| Conditioning - Yoke Control | BL Acq. | -0.1 | 53 | -0.4 | 0.692 |  |
| Conditioning - Yoke Control | BL Same Space | 0.1 | 53 | 0.4 | 0.724 |  |
| Conditioning - Yoke Control | BL Same Mov. | 0.2 | 53 | 1.1 | 0.294 |  |
| Conditioning - Yoke Control | Acq. | 1.5 | 53 | 5.1 | <0.001 | *** |

**Figure 3.**
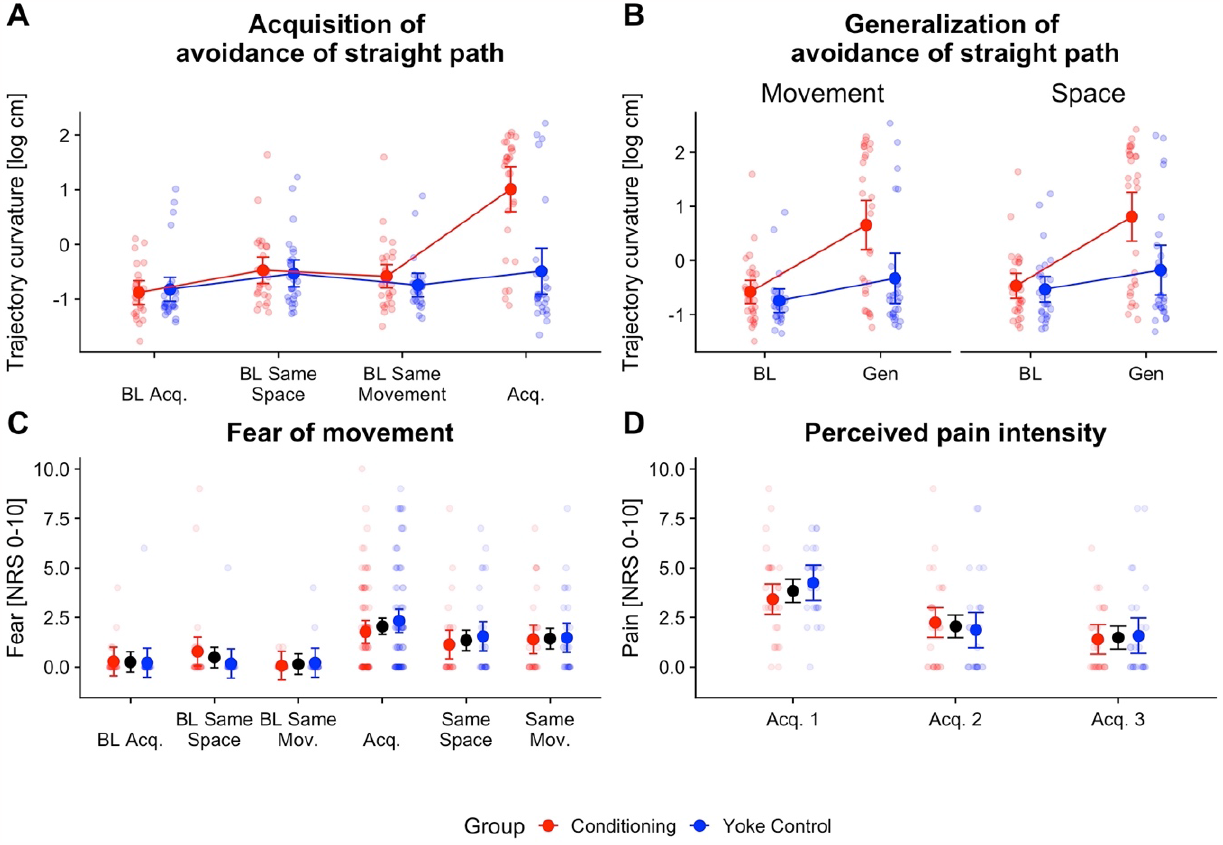
Raw data and statistical results. Raw data, averaged by participant, are shown as transparent points; the solid points are estimated marginal means resulting from the linear mixed models. Error bars indicate 95% confidence intervals. (A) Acquisition of trajectory curvature. Curvature was higher in the *Acquisition* phase than during baseline specifically in the *Conditioning* group (see text for details). (B) Comparison of trajectory curvature between generalization and baseline phases. Curvature was higher during generalization than baseline specifically in the *Conditioning* group, but results did not differ significantly between the types of association (space vs. movement). (C) Ratings of movement-related fear. Fear was higher during and after the *Acquisition* than during the baseline phases but did not differ between groups. (D) Ratings of pain intensity in the three *Acquisition* blocks. Perceived pain intensity declined over the blocks, but the decline did not depend on *Group*. In C and D, where *Group* did not play a significant role, black symbols represent the two groups combined.

The observed increase in trajectory curvature was consistent, observed in 26 out of 28 participants in the *Conditioning* group, and the effect size was large (Cohen’s d = 1.85).

### 3.2 Generalization of avoidance behavior and its association with space vs. movement

Generalization, i.e., the difference between *Baseline* and *Generalization* phases, depended on group, which was evident from a *Group* × *Phase* interaction in the LMM (F(1, 53) = 8.8, p = 0.005). Curvature was significantly stronger in *Generalization* than in *Baseline* phases in the *Conditioning* group (Diff_Gen-BL_ = 1.3 log cm, t(53) = 6.1, p < 0.001), but there was only a statistical trend in the *Yoke Control* group (Diff_Gen-BL_ = 0.4 log cm, t(53) = 1.8, p = 0.075), suggesting that participants of the *Conditioning* group, but not those of the *Yoke Control* group, responded to pain stimulation by systematically modifying their movements. Accordingly, post-hoc testing showed that curvature was significantly stronger in the *Conditioning* than the in the *Yoke Control* group in the *Generalization* phases (Diff_Cond-YK_ = 1.0 log cm, t(53) = 3.1, p = 0.003) but not in the baseline phases (Diff_Cond-YK_ = 0.1 log cm, t(53) = 0.8, p = 0.453); see Figure 3B). Notably, the factor *Association*, i.e. generalization based on movement vs. on space, did not interact with any other factor (see Table 4), indicating that the increase in curvature from *Baseline* to the *Generalization* phases was similar for the *Same Space* and *Same Movement* phases. There was, however, a main effect of *Association* (F(1, 53) = 8.4, p = 0.005): curvature was generally stronger in the phases designed to test spatial association (*BL Same Space* and *Same Space* combined) than in those designed to test movement association (*BL Same Movement* and *Same Movement* combined; Diff_Mov-Space_ = -0.2 log cm, t(53.3) = -2.9, p = 0.005). Note, this result is a main effect and, thus, affects *Baseline* and *Generalization* phases alike. Thus, this difference between spatial and movement association reflects differences in the overall characteristics of the movements required in the two contexts – for instance because they had different movement directions and angles – but was not related to the experimental manipulation of introducing pain stimuli. In sum, the model showed that generalization took place both for movement and space contexts to a similar extent; moreover, as one would expect, generalization was present only in the *Conditioning* but not the *Yoke Control* group.

**Table 4:** Results of the linear mixed model analyzing the increase in trajectory curvature from baseline to generalization phases between the groups.

| Term | num DF | den DF | F | p | sign. |
| --- | --- | --- | --- | --- | --- |
| Group | 1 | 53.0 | 7.7 | 0.007 | ** |
| Phase | 1 | 53.0 | 30.9 | <0.001 | *** |
| Association | 1 | 53.3 | 8.4 | 0.005 | ** |
| Group × Phase | 1 | 53.0 | 8.8 | 0.005 | ** |
| Group × Association | 1 | 53.3 | 0.2 | 0.646 | |
| Condition × Association | 1 | 4,732.1 | 0.1 | 0.815 | |
| Group × Phase × Association | 1 | 4,732.1 | 1.6 | 0.2 | |

**Table 4:** Pairwise-comparisons following the linear mixed model comparing the fear ratings between the groups and phases.

| contrast | estimate | DF | t | p | sign. |
| --- | --- | --- | --- | --- | --- |
| BL Acq. - Acq. | -1.8 | 375 | -7.9 | <0.001 | *** |
| BL Same Space - BL Same Mov. | 0.3 | 375 | 1.2 | 0.28 |  |
| BL Same Space - Acq. | -1.6 | 375 | -6.9 | <0.001 | *** |
| BL Same Space - Same Space | -0.9 | 375 | -3.1 | 0.003 | ** |
| BL Same Mov. - Acq. | -1.9 | 375 | -8.4 | <0.001 | *** |
| BL Same Mov. - Same Mov. | -1.3 | 375 | -4.6 | <0.001 | *** |
| Same Space - Same Mov. | -0.1 | 375 | -0.3 | 0.752 |  |

The fact that space and movement are both associated with pain on the group level could result from several patterns of individuals’ behavior. For instance, some participants may generalize based on space, and some based on movement, resulting in similar group averages for the two contexts. In such a scenario, curvature should correlate negatively between the two phases, as a given participant should express high avoidance in one, but not the other context. In contrast to this conjecture, the correlational analysis revealed a strong positive relationship between curvature in the *Same Space* and *Same Movement* phases for both groups (*Conditioning* group: r = 0.93, p < 0.001; *Yoke Control* group: r = 0.90, p < 0.001; see Figure 4). In other words, participants varied according to their overall tendency to generalize, and this tendency expressed itself in both generalization contexts and across both experimental groups.

**Figure 4.**
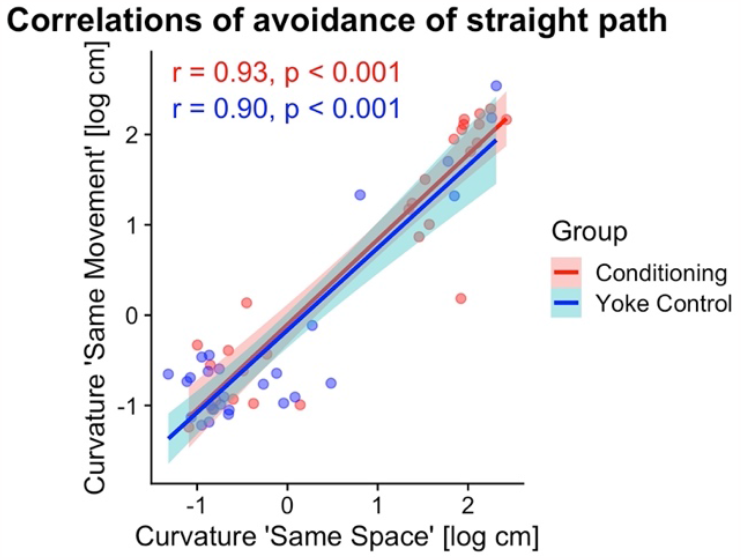
Correlation across generalization contexts, i.e., between trajectory curvature in the *Same Space* and the *Same Movement* phases. Avoidance was highly correlated across the two phases (see annotated Pearson correlations).

### 3.3 Fear and pain ratings

The LMM testing the differences in the fear ratings showed a significant main effect of *Phase* (F(5, 375) = 24.7, p < 0.001) but neither a main effect of *Group* (F(1, 56) = 0.1, p = 0.81) nor a significant *Group* × *Phase* interaction (F(5, 375) = 1.5, p = 0.19). This means that fear significantly differed between *Baseline* and *Acquisition*, but these differences did not depend on whether pain administration was contingent on one’s own behavior. Fear ratings were higher during *Acquisition* than in any baseline phase (see Table 4), and higher in both the *Same Space* (Diff_BL SS-SS_ = -0.9 log cm, t(375) = -3.1, p = 0.003) and Same *Movement* (Diff_BL SM-SM_ = -1.3 log cm, t(375) = -4.6, p < 0.001) generalization phases when compared to their respective baseline phases. Fear did not significantly differ between the *Same Space* and the *Same Movement* phases (Diff_SS-SM_ = -0.1 log cm, t(375) = -0.3, p = 0.752). The LMM comparing the groups’ pain intensity ratings indicated a main effect of *Block* (F(2, 110) = 29.8, p < 0.001), reflecting a slight decrease of perceived pain intensity over the three blocks of the *Acquisition* phase (see Figure 3D). No other effects were significant.

## 4 Discussion

In this study, we induced pain-associated avoidance behavior in a reaching task to investigate which kind of concurrent information becomes associated with the painful event: the movement itself, such as the kinesthetic cues related to movement direction, or the spatial location at which the pain was experienced – an aspect not addressed by previous studies (Glogan et al., 2020, 2022; Meulders et al., 2016). In medical contexts, it often seems sensible to assume that fear of pain is related to a movement, such as bending over in the context of back pain; yet one can imagine many other contexts in which pain is related to a location in space, such as when one is stung trying to cut a rose; in the latter case, pain is not related to the particular movement. Our concern was that experiments may try to test the former but actually (co-)implement the latter. Therefore, we disambiguated the two overlapping cues by employing two generalization contexts: one that required participants to make new movements through the spatial location at which they had previously received painful stimuli and one that required them to make the same movement as when pain had been applied, but within a different portion of space. We reasoned that avoidance should be strongest in the generalization context that shares the feature with the acquisition context that becomes associated with pain. Significant differences in avoidance and generalization behavior of the two experimental groups confirmed that our modified experimental paradigm adequately induced pain learning, replicating findings of previous studies (Glogan et al., 2020, 2022; Meulders et al., 2016) using a new movement paradigm.

### 4.1 Avoidance of both visuospatial and kinesthetic cues

The key finding of our study is that participants showed generalization of pain learning both with respect to the particular movement during which pain had previously been applied as well as with respect to the location at which the hand had been when the pain occurred. Hence, participants seemed to have inferred from the learning phase that they could evade pain by either steering clear of the critical spatial location or by avoiding specific movements.

The magnitude of avoidance was comparable between the generalization phases. This suggests that both types of cues, visuospatial and kinesthetic, were similarly involved in avoidance learning. Fitting with this result, there was a high correlation of avoidance over the two generalization contexts: when participants learned to avoid, they avoided both available (and confounded) cues. Given that the two aspects were (intentionally) confounded and ambiguous during the *Acquisition* phase this learning result is straightforward and in line with results from pioneering animal studies on classical conditioning with compound stimuli. Conditioning with a compound stimulus consisting of two CS that are paired with one US, for example a tone and a light followed by an electric shock, leads to the CR when one component of the compound is presented in isolation (Domjan, 2010; Kamin, 1969). In the present study, the kinesthetic and the visuospatial cues can be considered as parts of the conditioned compound and elicit the avoidance response when presented in isolation in the *Same Space* and *Same Movement* generalization conditions, respectively.

This could be explained with distinct associations being formed; the elements of the compound and the degree of avoidance are then an expression of the overlap of the learning and generalization contexts. During either of our generalization conditions, there is overlap with respect to only one of the two components of the compound. Accordingly, the degree of avoidance should be smaller during generalization than during learning, which is indeed what we observed in the present study. The notion of overlapping associations receives support from a previous study that separated the effects of movement endpoint (i.e., spatial cues) from movement direction (i.e., kinesthetic cues) on fear of movements (Meulders & Vlaeyen, 2019). In that study, participants of a *location* group received pain when they moved a joystick from the center to one side (CS+), but not when they moved it to the other side (CS-); participants of a *movement* group received pain when they moved the joystick from one side towards the center position (CS+) but not when they moved it from the other side (CS-). Notably, space and movement were effectively confounded, or overlapped, in that study’s *location* group, just like they were in the present study, because pain was presented when one defined movement was performed that ended in one specific location. In contrast, in that study’s *movement* group, pain was predicted only by movement, but not by location, because both CS+ and CS-movements shared the same center position endpoint. Fear of movements was stronger in the *location* group, consistent with the interpretation that fear will be higher the more features overlap with the learning context.

Our results are partly at odds with the fear-avoidance model of pain (Vlaeyen & Linton, 2000) because the model emphasizes the role of movement-pain associations but does not explicitly take into account visual-spatial information as relevant cues. Our findings, however, suggest that there may not be anything special about pain-movement coupling; instead, our results show that motor avoidance can equally well be associated with visuospatial cues and, in extension, it seems entirely sensible to suggest that it may rely on other types of cues in the same way.

It is, however, possible that learning processes are altered in chronic pain patients, which cannot be judged based on the current data of healthy controls. One possibility is that chronic pain patients may exhibit an increased tendency to associate pain with movement compared to other people, and that this bias promotes the emergence and/or maintenance of the vicious pain circle that is at the fear-avoidance model’s core. Alternatively, some evidence has suggested that chronic pain patients may exhibit a generally higher “conditionability”, i.e., a tendency toward faster learning via aversive conditioning mechanisms (Klinger et al., 2010; Schneider et al., 2004). Similarly, a potential role of altered sensory encoding and learned associations between pain and sensory input has been proposed elsewhere (Moseley & Vlaeyen, 2015; but see Fuchs, Becker, et al., 2015 for a critical review).

Our findings bear relevance for the conclusions drawn in previous studies that have used similar paradigms as the one employed here (Glogan et al., 2020; Meulders et al., 2016). These studies argued that the employed paradigm establishes fear and avoidance of specific movements; yet, pain may instead have been associated with the visual position of the hand cursor moving through specific spatial locations. On the one hand, our present results support the conclusion drawn by the discussed studies that a pain-movement association has been formed; on the other hand, they suggest that those previous studies may have overlooked associations formed between pain and visual-spatial cues. In fact, we would argue that visuospatial associations were likely even stronger in their paradigm than in ours: in those studies, both the hand cursor and the relevant spatial locations (gates) were presented visually. In our study, the relevant location was not visualized. Thus, the interpretation of previous findings as implying movement-based associations may be correct but incomplete.

Yet, one of the discussed previous findings may, in fact, indicate that the previously employed paradigm predominately induced space-based associations. Recall that no generalization of avoidance was observed in the study by Glogan et al. (2020) when the visual layout of the experimental scene was modified after the learning phase by changing the layout of the gates through which participants had to guide the visual cursor. This result can be explained by assuming that the learned association was based on visual cues and that participants could easily discriminate new from old gates visually. Thus, the perceived similarity between the gates was low and presumably prevented generalization of avoidance. As the authors themselves argue in their article, curiosity might have motivated participants to explore the new location, resulting in rapid extinction of avoidance behavior. Whether or not our suggestion turns out to be true—it demonstrates that disregarding the distinction between space and movement in experimental paradigms aimed at investigating pain may lead to misinterpretations of experimental findings and highlights that adaptations to the currently established paradigms are called for. This is not to say that space-related learning must necessarily be key for how patients develop their vicious pain circles. Our argument is rather that paradigms targeted at pain-movement learning may actually be testing something categorically different. Polemically put, we may intend to design an experiment analogous to patients learning to avoid back bending, but accidentally implement a paradigm that is analogous to cutting a thorny rose. Scientifically speaking, we must design experiments that investigate pain-movement associations in ways that rule out alternative explanations based on visual and spatial learning instead.

### 4.2 Similar pain-induced fear of movement in both groups

As predicted, the introduction of painful stimuli led to an increase in movement-related fear. This increase was evident not only in the *Conditioning* group, but also in the *Yoke Control* group, which had no control over stimulus occurrence and intensity. In fact, the general levels of movement-related fear were comparable between the two groups, which is compatible with the overall fear levels being similar for experimental and yoked participants in the study in which a cursor was moved through gates (Meulders et al., 2016). However. in that study, participants of the experimental group feared movements through the pain-associated gate most, whereas yoked participants feared all gates alike. We cannot differentiate movements at that level in the present study, because participants chose their trajectories freely and we assessed fear levels as a ratings after full blocks, which accordingly comprised straight and curved trajectories.

### 4.3 Conclusions, outlook, and open questions

In summary, our study showed that visuospatial and kinesthetic information are associated with pain when the respective cues are available simultaneously and their role for the occurrence of pain is ambiguous. This shows that humans exploit visuospatial information in addition to kinesthetic cues for pain learning, an aspect that has been neglected both by the fear-avoidance model of pain and by most previous research (with the exception of Meulders & Vlaeyen, 2019). Interpretations of previous findings may therefore be incorrect or, at least, incomplete. For now, it remains an open question whether spatial and kinesthetic information can, in principle, be separated and, for instance, be integrated with individual weights during pain learning. Clearly, however, findings from the motor control literature, both regarding the different timing for proprioceptive, visual effector, and visual target-related processing (Scott, 2016), as well as regarding learning principles, make this a sensible hypothesis (Avraham et al., 2022; Franklin & Wolpert, 2011).

Similarly, the clinical relevance of visuospatial associations, as identified here, remains to be determined. With our example of reaching for a thorny rose, we suggested that current pain paradigms may unintentionally test the role of the rosebush (i.e., the rose’s location) instead of the reach (i.e., the movement). Our results imply that both space-based and movement-based associations are established concurrently, with neither dominating the other. However, when patients experience acute pain that sets the fear and avoidance circle in motion, proprioceptive cues might play a more important role.

Some studies have demonstrated that chronic pain patients exhibit deficits in proprioception, for example when they estimate the position of their limbs in space (Brun et al., 2019). Impaired proprioception might also relate to altered movement-related learning processes such as exaggerated generalization of fear and avoidance (Vandael et al., 2022). It is a common assumption in the learning literature on stimulus generalization that worse discrimination between stimuli is associated with a wider generalization gradient (Guttman & Kalish, 1956; Honig & Urcuioli, 1981). This concept is related to a more recent proposal termed the “imprecision hypothesis” that highlights the role of sensory imprecision in learning and chronic pain (Moseley & Vlaeyen, 2015). So, is the visuospatial conditioning that the laboratory paradigms (including ours) invoke a relevant aspect of pain-induced learning outside the lab, or is it rather a nuisance factor that limits the ecological validity of our current paradigms?

Some evidence suggests that key aspects in the learning of movement avoidance in response to pain might not be kinesthesis but rather posture. Accordingly, all sensory modalities that contribute to posture perception might then contribute to maladaptive pain learning. First, several studies have reported that patients exhibit pain-related fear not only to movements but also to specific postures shown on static images (Gatzounis et al., 2021; Meulders et al., 2015, 2017). Second, neck pain patients expect pain for certain movement angles of their head, but their expectation of pain can be modulated by providing false visual feedback about head rotation in virtual reality (Harvie et al., 2015).

### 4.4 Summary

In summary, our research indicates that the learning processes in simplified laboratory settings are more intricate than previously accounted for, revealing that participants form associations with multiple available cues. When this laboratory observation is extrapolated and applied to a natural, less controlled scenario with multiple stimuli available simultaneously, it is plausible that chronic pain-related associations could extend beyond just movement and relate to other factors. These factors could include other sensory cues, such as proprioception and vision, as well as contextual cues related to the surrounding environment or social context. Movement and posture may have a significant impact on chronic pain. However, it is critical to accurately identify the underlying associations and provide clean evidence to support this claim. A close examination of the role of different sensory modalities involved in pain-related learning processes might be a promising approach to shed light on underlying processes that underlie pain learning and might help translating research into successful treatment of chronic pain.

## Acknowledgements

We thank Evelyn Schulz for assistance with the data acquisition.

## Conflicts of interest

The authors declare that there was no conflict of interest.

## Notes

https://osf.io/rjz36/

### Competing Interest Statement

The authors have declared no competing interest.

### Summary of Updates

The abstract has been shortened

