## Supplementary Material for "The role of kinesthetic and visuospatial cues in pain-induced movement avoidance"

#### 1 Statistical models with and without log-transformation

We used log-transformed curvature values for the linear mixed models (LMM) because the distribution of the (untransformed) values was less suitable for the statistical models. Here we show that the LMM applied to compare the baseline phases with the acquisition phase, reported in section 3.1 in the main text, fits the data better when a log-transformation is applied, but that conclusions drawn when using untransformed values remain the same.

##### 1.1 Distribution of curvature values and model residuals

Figure S1 shows the distribution (histogram) of the untransformed (left) and the log-transformed (right) dependent variable. As can be seen in Figure S1, the untransformed curvature value's distribution has many observations at the lower end and a long tail to the right side (with high values). The log-transformed variable shows a slight bimodality, but the transformation clearly reduces the potential problem of a long tail on the right side.

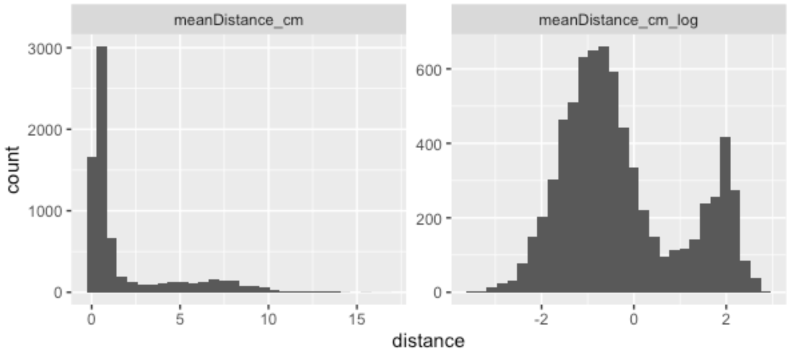


Figure S1: Histograms of the untransformed (left) and the transformed (right) curvature values.

That the distribution of the untransformed data leads to a problem when entering the LMM is shown by the distribution of the LMM's residuals, shown in Figure S2 A. The distribution has heavy tails on both sides, violating normality of residuals. This problem is drastically reduced when using the log-transformed variable, as shown in Figure S2 B.

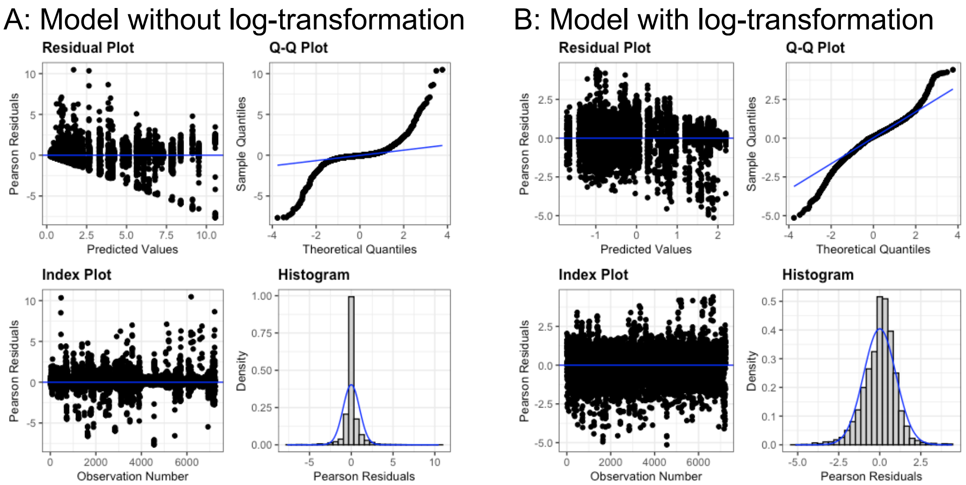

Figure S2: Residuals of the LMM based on the untransformed dependent variable (A) and based on the log-transformed dependent variable.

#### 1.2 Comparability of statistical results

Comparing Table S1 to Table 1 in the main text makes obvious that statistical results are not substantially affected by the transformation and the conclusions remain the same.

| Term | num DF | den DF | F | p | sign. |
|---|---|---|---|---|---|
| Group | 1 | 53.0 | 7.2 | 0.01 | ** |
| Phase | 3 | 52.9 | 16.9 | <0.001 | *** |
| Group:Phase | 3 | 52.9 | 5.6 | 0.002 | *** |

Table S1: Main effects and interactions of the LMM based on the untransformed dependent variable.

### 2 Sensitivity analysis

#### 2.1 Sensitivity of the acquisition of avoidance effect

All details about the sensitivity analysis are provided in the reproducible R-code publicly available in the accompanying OSF repository: https://osf.io/rjz36/

The sensitivity analyses comprised two parts. In the first part, we tested how much the effect size between the baseline phases and the *Acquisition* phase in the *Conditioning* group can be decreased before the LMM does not indicate the significant effects any longer. The hypotheses tested in the simulations were:

- The critical test is that the difference between the baseline and the *Acquisition* phase needs to be larger in the conditioning than in the yoke control group. This is captured by the *Phase x Group* interaction (Hypothesis 1)
- In addition, the curvature values in the *Acquisition* phase also need to be significantly larger than at baseline for the *Conditioning* group. This is expressed by a significant negative difference value (baselines - *Acquisition*) in the post-hoc test for the *Conditioning* group (Hypothesis 2)

We decreased the effect size in 10% steps and repeated the LMM for each level. To additionally determine the role of the sample size, we also iteratively left out more and more participants (chosen at random) and repeated the analysis with 250 iterations per combination of effect size and number of participants.

The results are shown in Figure S3.

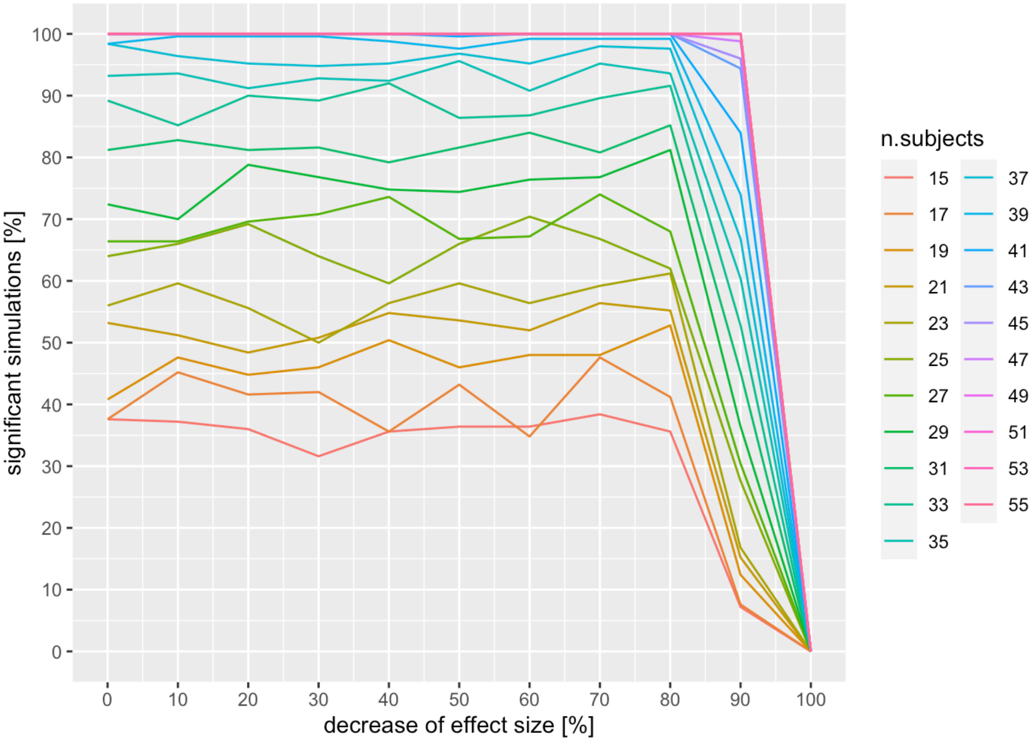


Figure S3. Results of the sensitivity analysis of the acquisition of avoidance in the conditioning group.

Figure S3 shows that the study was very well powered to detect a difference between the groups in the difference between baseline and the *Acquisition* phase. In fact, the effect was still found when decreasing the effect size down to as much as 90% of the original effect size. When using fewer participants, the effect became less robust. We conclude that the study with the present sample was well powered to detect effects even if they had been considerably smaller than the observed effects. However, assessing fewer participants in future studies will considerably increase the risk of a type 2 error.

#### 2.2 Sensitivity of the generalization of avoidance effect

We took the same approach to test specific increased generalization in the *conditioning* group. In analogy, the hypotheses were:

- A significant *Phase x Group* interaction (Hypothesis 1)
- A significant negative difference value (baseline - *Generalization*) in the post-hoc test for the *Conditioning* group (Hypothesis 2)

The results are shown in Figure S4.

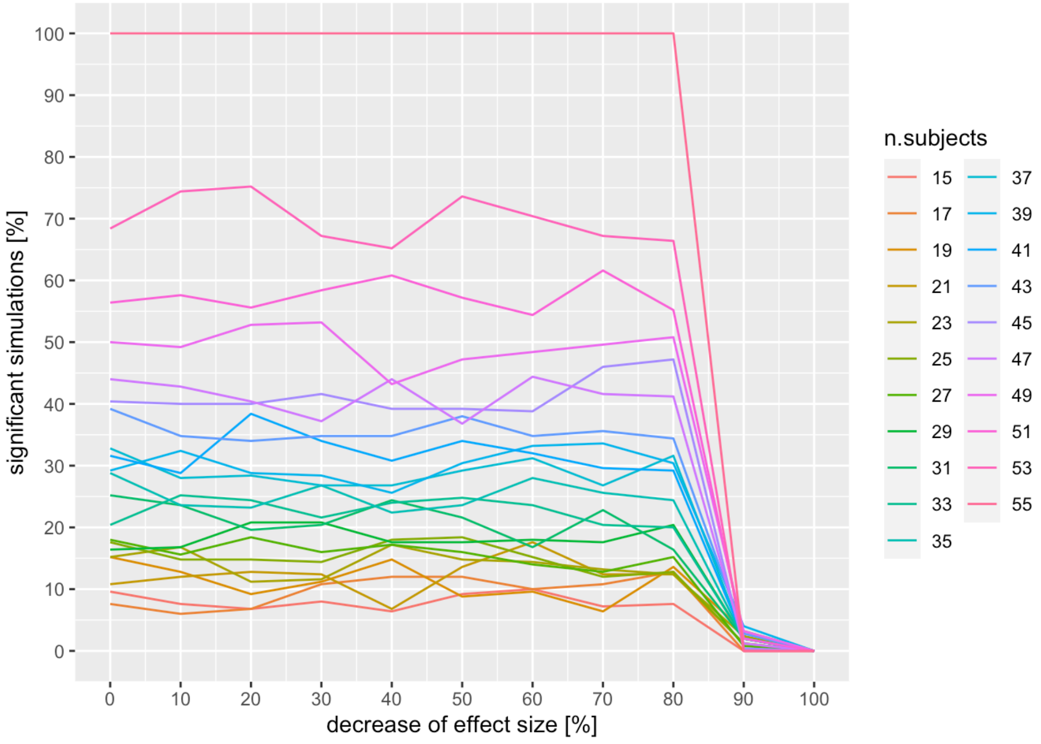


Figure S4. Results of the sensitivity analysis of the generalization of avoidance in the conditioning group.

Figure S4 shows that the study was very well powered to detect a difference between the groups in the difference between baseline and generalization phases. The effect was still found when decreasing the effect size down to as much as 80% of the original effect size. When using fewer participants, the effect became less robust. As before, we conclude that the study with the present sample was well powered to detect effects even if they had been considerably smaller than the observed effects but that fewer participants in future studies will considerably increase the risk of a type 2 error.
